# Differences in immune cell profiles around the time of islet autoimmunity seroconversion in children with and without type 1 diabetes

**DOI:** 10.1101/2025.06.23.661117

**Authors:** Kirk R. Hohsfield, Patrick M. Carry, Sarah D. Slack, Charlie T. Repaci, Lauren A. Vanderlinden, Katerina Kechris, Marian Rewers, Jill M. Norris, Randi K. Johnson

**Author notes:** Corresponding author: Randi K. Johnson. Email addresses.

## Abstract

Seroconversion (SV) marks the initiation of islet autoimmunity (IA) and pre-clinical phase of type 1 diabetes, yet the contributions of immune cells beyond cytotoxic T cells remain unclear. We applied high-resolution immune cell-type deconvolution using peripheral blood DNA methylation data from nested case-control samples of the Diabetes Autoimmunity Study in the Young (DAISY; n=151) and The Environmental Determinants of Diabetes in the Young (TEDDY; n=166) to estimate immune cell proportions across pre-SV and SV timepoints and construct functional ratios, such as the neutrophil-to-lymphocyte ratio (NLR). Using linear models, we evaluated differences between type 1 diabetes cases and controls at pre-SV, SV, and the change across timepoints. Pre-SV, cases had higher NLR and lower CD4T/CD8T cell ratios. At SV, the combined B-CD4T-CD8T memory/naïve ratio was reduced in cases. From pre-SV to SV, cases showed attenuations in NLR, B-memory/naïve, and B-CD4T-CD8T memory/naïve ratios. These patterns may reflect delayed or disrupted immune maturation with the persistence or expansion of naïve cells or impaired transition to memory subsets following antigen exposure. Our findings highlight early shifts in innate and adaptive immune cell dynamics during type 1 diabetes pathogenesis and support immune cell ratios as potential biomarkers for risk stratification and mechanistic insight.

**Article Highlights:**

- We sought to examine immune cells around the time of IA seroconversion in children at higher risk for type 1 diabetes.
- We wanted to answer whether immune cell ratio differences exist between type 1 diabetes cases and controls around IA at pre-SV, SV, and the change pre-SV to SV.
- We found immune cell ratio differences between type 1 diabetes cases and controls before, during, and across SV timepoints, suggesting potential etiological and pathophysiological roles.
- Our findings highlight the complexity of immunodynamics around IA seroconversion and potential role for immune cell ratios in type 1 diabetes risk stratification and intervention.

## Introduction

Type 1 diabetes remains one of the most common chronic autoimmune diseases affecting children globally[1]. Characterized by immune-mediated destruction of insulin-producing β-cells in the pancreas, it ultimately leads to hyperglycemia, ketoacidosis, and a lifelong dependency on exogenous insulin therapy [1]. Current global estimates point to increasing incidence rates, especially among children and adolescents, with estimated average annual increases of approximately 3-4% over the last three decades, highlighting a significant public health burden [2–4]. Prior to developing type 1 diabetes, islet autoimmunity (IA) occurs, defined by the presence of autoantibodies, such as insulin (IAA), glutamic acid decarboxylase (GADA), insulinoma antigen-2 (IA-2A), and zinc transporter 8 (ZnT8A), targeting pancreatic islet cells. IA signifies early faltering in immune tolerance and marks the beginning of a variable progression towards type 1 diabetes, although some people do not progress to disease [5, 6]. Despite established associations between IA and type 1 diabetes, gaps remain in our understanding of the underlying immune processes.

Immune cell dynamics that occur around the time of IA seroconversion (SV) and during progression from IA to type 1 diabetes have not been fully elucidated. Recent advances in immunophenotyping have emphasized the utility of peripheral blood-based immune cell ratios – such as neutrophil-to-lymphocyte (NLR), CD4T/CD8T ratio, and memory/naïve (mem/nv) lymphocyte ratios – as potential biomarkers of systemic inflammation, immune activation, and immune maturation status [7, 8]. Specifically, the NLR indicates systemic inflammation and adaptive immunity, the CD4T/CD8T ratio reflects immune senescence and activation, and the B and T-cell mem/nv ratios reveal levels of balance between the cells of the adaptive immune system [7–10]. From these ratios, we can glean insight into the balance between immune cell subsets rather than focusing on absolute or relative abundance alone, which can help to characterize underlying immune dynamics, such as shifts between innate and adaptive immune responses or naïve and antigen-experienced states. Differential ratios of immune cell populations, therefore, can reflect underlying disease processes and may predict onset or worsening of disease. Ratios offer modeling options that address the inherent compositional nature of peripheral blood-based cell proportion data (i.e., the sum-to-one constraint, whereby a change in one cell proportion affects the others), thereby capturing biologically meaningful contrasts while avoiding potential spurious associations driven by interdependence among components [11, 12]. However, few studies have utilized longitudinal data to examine changes in immune cell ratios before and after the loss of immune tolerance associated with the onset of IA in pediatric populations.

This study aims to bridge these knowledge gaps by evaluating immune cell type heterogeneity and dynamics during critical time windows before and after IA seroconversion. Leveraging two well-established prospective cohorts — the Diabetes Autoimmunity Study in the Young (DAISY) and The Environmental Determinants of Diabetes in the Young (TEDDY) — we estimated immune cell ratios derived from DNA methylation (DNAm) in peripheral blood to better understand the immunological shifts occurring in children genetically susceptible for type 1 diabetes during a critical stage of disease pathogenesis.

## Research Design and Methods

### DAISY study population

The prospective DAISY cohort enrolled and followed 2,547 children based around Denver, Colorado. They were recruited either as part of population-based newborn screening at St. Joseph’s Hospital for specific human leukocyte antigen (HLA) genotypes, known to have increased genetic risk of type 1 diabetes, or from unaffected first-degree relatives of current patients, then followed for the development of IA and type 1 diabetes. Additional study information is detailed elsewhere [13]. Clinic visits with blood sample collection occurred at 9, 15, and 24 months of age, then annually thereafter until the development of IA. Participants who developed IA came in for visits every 3-6 months until type 1 diabetes diagnosis, based on American Diabetes Association (ADA) clinical guidelines. Most participants identified as Non-Hispanic White among the race-ethnicity categories. The study protocol was approved by the Colorado Multiple Institutional Review Board (COMIRB) and adhered to all relevant research regulations. Informed consent was obtained by the parents or primary caretakers of all participating children, and assent was obtained from children seven years of age and older.

A nested case-control subset of DAISY study participants was selected to evaluate epigenetics – specifically, DNAm – and other omics measurements in the context of IA and type 1 diabetes. Type 1 diabetes cases were frequency matched on age at IA seroconversion, race-ethnicity, and sample availability across five timepoints – birth, infancy (age 9-16 months), pre-IA, onset of IA, and prior to diagnosis of type 1 diabetes. This analysis included type 1 diabetes cases and controls with good quality DNAm data available pre-IA, at IA onset, or both (n = 151). DAISY defined IA as the persistent presence of one or more autoantibodies – IAA, GADA, IA-2A, and ZnT8A – detected across two consecutive visits, or as a single positive autoantibody result followed by a type 1 diabetes diagnosis at the next visit [14–16].

### TEDDY study population

TEDDY represents a multi-national prospective cohort study of children followed from birth with an increased genetic risk of type 1 diabetes, based on their HLA genotype, and included 8,676 participants that were followed through the development of IA until type 1 diabetes diagnosis. More specific study design and follow-up details are described elsewhere [17, 18]. Recruitment occurred across six clinical research centers: three in the USA (Colorado, Georgia/Florida, and Washington) and three in Europe (Germany, Finland, and Sweden). Visits were conducted every three months until age 4, then every six months until age 15. Similar to DAISY, an accelerated schedule was used after the development of IA, which included visits every three months until age 15 years or diagnosis of type 1 diabetes based on ADA clinical criteria. The study protocol was approved by local Institutional Review Boards for each of the six clinical centers and adhered to all relevant research regulations. An external review committee formed by the NIH monitored the study. Informed consent was obtained by the parents or primary caretakers of all participating children, and assent was obtained from children seven years of age and older.

From the TEDDY cohort, a type 1 diabetes risk-set based on a nested case-control cohort with DNAm data was selected. Type 1 diabetes cases were matched 1:1 with controls based on age, sex, type 1 diabetes family history, and clinical center. Study participants are primarily composed of persons identifying as Non-Hispanic White, and a majority were recruited from the European clinical centers. We included type 1 diabetes cases where age at developing IA was known and who had good quality DNAm data available pre-IA, onset of IA, or both, along with samples from matched controls (n = 166). Similar to DAISY, TEDDY defined IA as testing positive for one of IAA, GADA, or IA-2A across two consecutive visits [19].

### Study timepoints

Although the DAISY and TEDDY studies prospectively collected multiple samples per participant, for this analysis, we selected one sample per subject at each available timepoint of interest: before IA (pre-SV) and at IA seroconversion (SV), when participants converted from seronegative to seropositive with the detection of islet autoantibodies. The pre-SV timepoints in DAISY and TEDDY included 84 and 122 samples, respectively. The SV timepoints in DAISY and TEDDY included 140 and 160 samples, respectively.

### DNA methylation

DNAm measures were obtained from whole blood samples on the Infinium HumanMethylation450 (450K) and MethylationEPIC (EPIC) BeadChip arrays (Illumina, San Diego, CA, USA), which interrogate over 450,000 and 850,000 methylation sites, respectively, across the genome at single-nucleotide resolution. DAISY participants had samples from both 450K and EPIC arrays while TEDDY participants only included samples from EPIC arrays. Samples of low quality or with discordant values for sex were removed. Technical replicate samples were included during preprocessing for quality control, as detailed in previous work in DAISY [20] and TEDDY [unpublished], and were removed prior to statistical analysis.

### Cell deconvolution

We performed cell deconvolution using the estimateCellCounts2 function within the FlowSorted.Blood.EPIC package (v2.8.0) and the cell references from the FlowSorted.BloodExtended.EPIC package (v1.1.2) in R, utilizing the recommended parameters for CpG selection and normalization [21, 22]. The latter included an expanded reference for 12 immune cell proportions: neutrophils (Neu), eosinophils (Eos), basophils (Baso), monocytes (Mono), naïve and memory B cells (Bnv, Bmem), naïve and memory CD4+ and CD8+ T cells (CD4Tnv, CD4Tmem, CD8Tnv, CD8Tmem), natural killer (NK), and T regulatory (Treg) cells. If any individuals had zero values in their cell proportions, we imputed them via substitution by taking the minimum value within each cell type then dividing it by two [23]. We used previously published immune cell references to gauge whether our participants were within expected ranges for certain subsets [24]. Compared to published immune cell references, TEDDY participants had lower immune cell proportions, although they overlapped between the 10^th^ and 90^th^ percentiles. **Supplemental Table 1** provides a snapshot of several immune cell subsets (B, CD4T, CD8T, and NK) for TEDDY samples collected between 1-2 years of age, which includes the average age before and during SV.Statistical approach

We tested whether immune cell ratios differed between type 1 diabetes cases and controls at pre-SV, SV, or changed differently from pre-SV to SV using linear modeling of immune cell ratios as the outcome and an interaction between type 1 diabetes status and time as the primary covariates of interest. For the outcomes, we used log-transformed immune cell ratios, which included the following: neutrophil/lymphocyte (NLR), monocyte/lymphocyte (MLR), B-memory/B-naïve (B-mem/nv), CD4T-memory/CD4T-naïve (CD4T-mem/nv), CD8T-memory/CD8T-naïve (CD8T-mem/nv), CD4T/CD8T, and B-CD4T-CD8T-memory/B-CD4T-CD8T-naïve (B-CD4T-CD8T-mem/nv). Lymphocytes consisted of CD4+ and CD8+ T cells, B memory and naïve cells, natural killer (NK) cells, and T regulatory cells.

Type 1 diabetes status (case vs. control) served as our primary explanatory variable to investigate immune cell ratio differences between these groups. A covariate for time was included to distinguish between the two different seroconversion measurements (pre-SV vs. SV). We included an interaction term between type 1 diabetes status and time to consider how immune cell ratios varied by type 1 diabetes status across two different sample collection timepoints. In DAISY, additional covariates for age, DR3/4 HLA status, and platform were included to either adjust for confounding or increase precision. For TEDDY, we included DR3/4 HLA status as a fixed effect and the case-control index variable (comprised of age, sex, type 1 diabetes family history, and clinical center) as a random intercept to account for the matching strata. Type 1 diabetes family history represented a dichotomous variable (yes vs. no). For reporting, we exponentiated each immune cell ratio, yielding the immune cell proportion ratio between type 1 diabetes cases and controls (geometric mean), and presented them as percentage differences. The lm base function in R and the lmer function from the lme4 package (v1.1-35.5) were applied to construct the linear and linear mixed-effects models, respectively [25]. We used an alpha level of 0.05 to determine statistical significance across all analyses.

We estimated the effects of interest separately by study. Then, given the similarities in study design (eligibility, follow-up, IA and type 1 diabetes estimation), we combined them with inverse variance-weighted, fixed effects meta-analysis using the meta package in R [26]. We evaluated heterogeneity based on Cochran’s Q and I^2^, following through with the meta-analysis in all instances of low estimated variability between study effects. To evaluate the robustness of our findings, we performed two sensitivity analyses – one that excluded samples with imputed cell proportion values and another which assessed differences by three-level type 1 diabetes family history (mom, dad/sibling, none). A combination of *a priori* knowledge, Akaike information criterion (AIC), and consistency of effect size and direction were examined to determine the most appropriate models.

### Data and Resource Availability

A subset of the DNAm data analyzed during the current study is available through NCBI GEO: GSE142512. Investigators may request access to additional data through the DAISY NIDDK repository: https://repository.niddk.nih.gov/study/206. DNAm data for TEDDY may be requested through an ancillary study (https://teddy.epi.usf.edu/research/). Additional data can be found in the TEDDY NIDDK repository: https://repository.niddk.nih.gov/study/24. All analyses were performed using R Statistical Software 4.4.0 (R Core Team, 2024) [27]. Code used in the statistical analysis will be available through the lab GitHub: https://github.com/rkjcollab. This research was enabled, in part, by the use of the FlowSorted.BloodExtended.EPIC software package developed at Dartmouth College, which is provided under a free academic license as specified here: https://github.com/immunomethylomics/FlowSorted.BloodExtended.EPIC/blob/main/SoftwareLicense.FlowSorted.BloodExtended.EPIC%20to%20sign.pdf. The software is subject to the licensing terms made available by Dartmouth Technology Transfer and is provided “AS IS” without any warranties, express or implied.

## Results

Participant characteristics for the DAISY and TEDDY cohorts are summarized in **Table 1**, along with sample sets and intersections displayed as UpSet plots in **Figure 1** [28]. Participants are distinguished by type 1 diabetes status, while the samples are further categorized across pre-SV and SV timepoints. In DAISY, a total of 151 children were included in the analysis, where 76 developed type 1 diabetes. Values for the frequency-matched variables (age at seroconversion, race/ethnicity, and sample availability) remained similar between groups. Type 1 diabetes cases had a higher proportion of first-degree relatives with type 1 diabetes as well as DR3/4 HLA genotypes compared to controls. Across the 224 participant samples in the DAISY analysis, there were 111 samples from type 1 diabetes cases and 113 from controls. These consisted of 84 and 140 samples for the pre-SV and SV time points, respectively, as illustrated in **Figure 1A**. In TEDDY, a total of 166 children were included for analysis with 83 developing type 1 diabetes. The matching variables (age, sex, type 1 diabetes family history, and clinical center) were closely aligned between groups. Type 1 diabetes cases had a higher proportion of DR3/4 HLA genotypes compared to controls. A total of 282 participant samples were included in the analysis, comprised of 141 samples from type 1 diabetes cases and 141 from matched controls. These were composed of 122 and 160 samples for pre-SV and SV time points, respectively, per **Figure 1B**. The average age at which DAISY cases developed type 1 diabetes was older than TEDDY at 9.48 years compared to 2.86 years, respectively.

**Figure 1.**
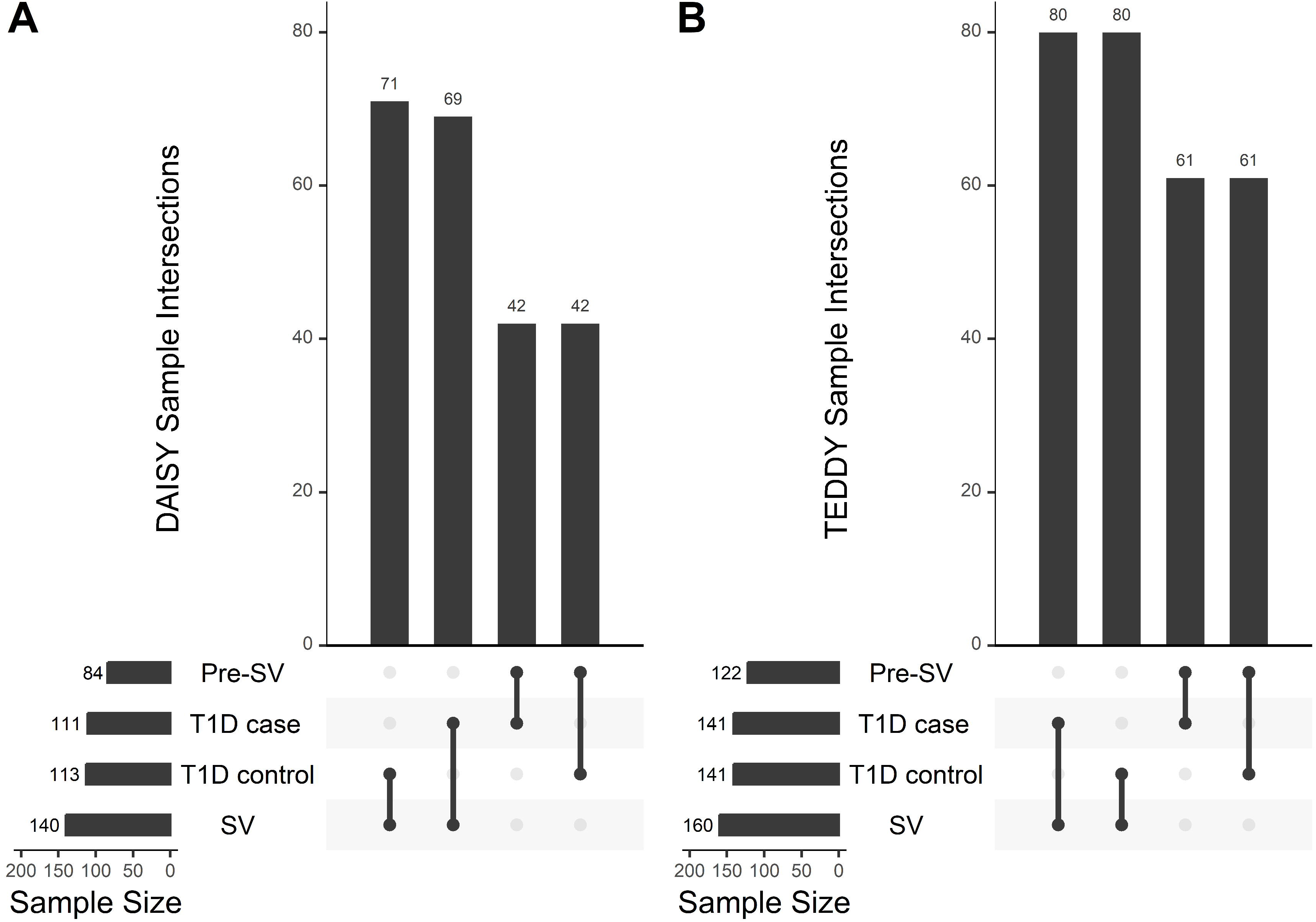
UpSet plots with sample breakdowns by study cohort. * Abbreviations: Pre-SV (pre-seroconversion), SV (seroconversion), T1D (type 1 diabetes). Figure 1A captures the different sample set sizes and intersections from DAISY participants while Figure 1B depicts samples from TEDDY participants. In both cohorts, more SV samples were available compared to Pre-SV samples, based on the quality and sample selection criteria for the present study (Research Design and Methods).

**Table 1.** Participant Characteristics for DAISY and TEDDY Participants.

|  | **<u>DAISY</u>** - - |  |  | **<u>TEDDY</u>** - - |  |  |
| --- | --- | --- | --- | --- | --- | --- |
|  | **T1D**<br>**Control**<br>**N=75** | **T1D Case**<br>**N=76** | **Total**<br>**N=151** | **T1D**<br>**Control**<br>**N=83** | **T1D Case**<br>**N=83** | **Total**<br>**N=166** |
|  | N (%) or Mean (SD) |  |  | N (%) or Mean (SD) |  |  |
| **Sex (female)** | 30 (40.0) | 37 (48.7) | 67 (44.4) | 36 (43.4) | 36 (43.4) | 72 (43.4) |
| **Non-Hispanic White (yes)** | 68 (90.7) | 68 (89.5) | 136 (90.1) | 57 (68.7) | 55 (66.3) | 112 (67.5) |
| **T1D family history (yes)** | 41 (54.7) | 54 (71.1) | 95 (62.9) | 27 (32.5) | 27 (32.5) | 54 (32.5) |
| **Age at IA onset (years)** | - | 3.78 (3.12) | - | - | 1.34 (0.72) | - |
| **Age at T1D onset (years)** | - | 9.48 (4.79) | - | - | 2.86 (1.37) | - |
| **HLA-DR3/4 (yes)** | 14 (18.7) | 34 (44.7) | 48 (31.8) | 32 (38.6) | 48 (57.8) | 80 (48.2) |
| **Platform (450K)** | 37 (49.3) | 38 (50.0) | 75 (49.7) | 0 (0) | 0 (0) | 0 (0) |
| **Clinical Center** |  |  |  |  |  |  |
| Georgia/Florida |  |  |  | 5 (6.0) | 5 (6.0) | 10 (6.0) |
| Washington |  |  |  | 5 (6.0) | 5 (6.0) | 10 (6.0) |
| Colorado | 75 (100) | 76 (100) | 151 (100) | 12 (14.5) | 12 (14.5) | 24 (14.5) |
| Sweden |  |  |  | 25 (30.1) | 25 (30.1) | 50 (30.1) |
| Germany |  |  |  | 10 (12.0) | 10 (12.0) | 20 (12.0) |
| Finland |  |  |  | 26 (31.3) | 26 (31.3) | 52 (31.3) |

**Figure 2** illustrates the results from the inverse variance-weighted fixed effects meta-analysis, which are summarized below. **Figure 2** captures the consistency between the effects of the DAISY and TEDDY studies, as well as the pooled effect. This approach enabled us to detect the B-mem/nv and B-CD4T-CD8T-mem/nv ratios that might have been overlooked if we had analyzed the studies independently. In the pre-SV period, the NLR was 15% higher (Ratio: 1.15, 95% CI: 1.00, 1.33; p = 0.0442) while the CD4T/CD8T was 9% lower (Ratio: 0.91, 95% CI: 0.83, 1.00; p = 0.0448) in type 1 diabetes cases compared to controls after adjusting for covariates. The remaining immune cell ratios were null for the pre-SV timepoint. In the SV period, the B-CD4T-CD8T-mem/nv ratio was estimated to be 26% lower (Ratio: 0.74, 95% CI: 0.57, 0.98; p = 0.0325) among type 1 diabetes cases compared to controls, after covariate adjustments. The other immune cell ratios during SV did not yield statistically significant effects. Results for the pre-SV to SV change are displayed across **Figure 2** and further detailed in **Figure 3**. For the pre-SV to SV change analysis, we found that the B-mem/nv, B-CD4T-CD8T-mem/nv, and NLR ratio trajectories were reduced by 35% (Ratio: 0.65, 95% CI: 0.43, 0.96; p = 0.0324), 38% (Ratio: 0.62, 95% CI: 0.41, 0.95; p = 0.0273), and 21% (Ratio: 0.79, 95% CI: 0.66, 0.95; p = 0.0119), respectively, among type 1 diabetes cases compared to controls, after adjusting for covariates. **Figure 2** reveals that type 1 diabetes cases started with higher immune cell ratios for the B-mem/nv, B-CD4T-CD8T-mem/nv, and NLR, but controls had more pronounced increases at seroconversion. No other significant associations were found for the additional immune cell ratios during the pre-SV to SV change. Results were robust to sensitivity analyses, and we did not observe immune cell ratio differences by type 1 diabetes family history, though, we were likely underpowered for the three-way interaction.

**Figure 2.**
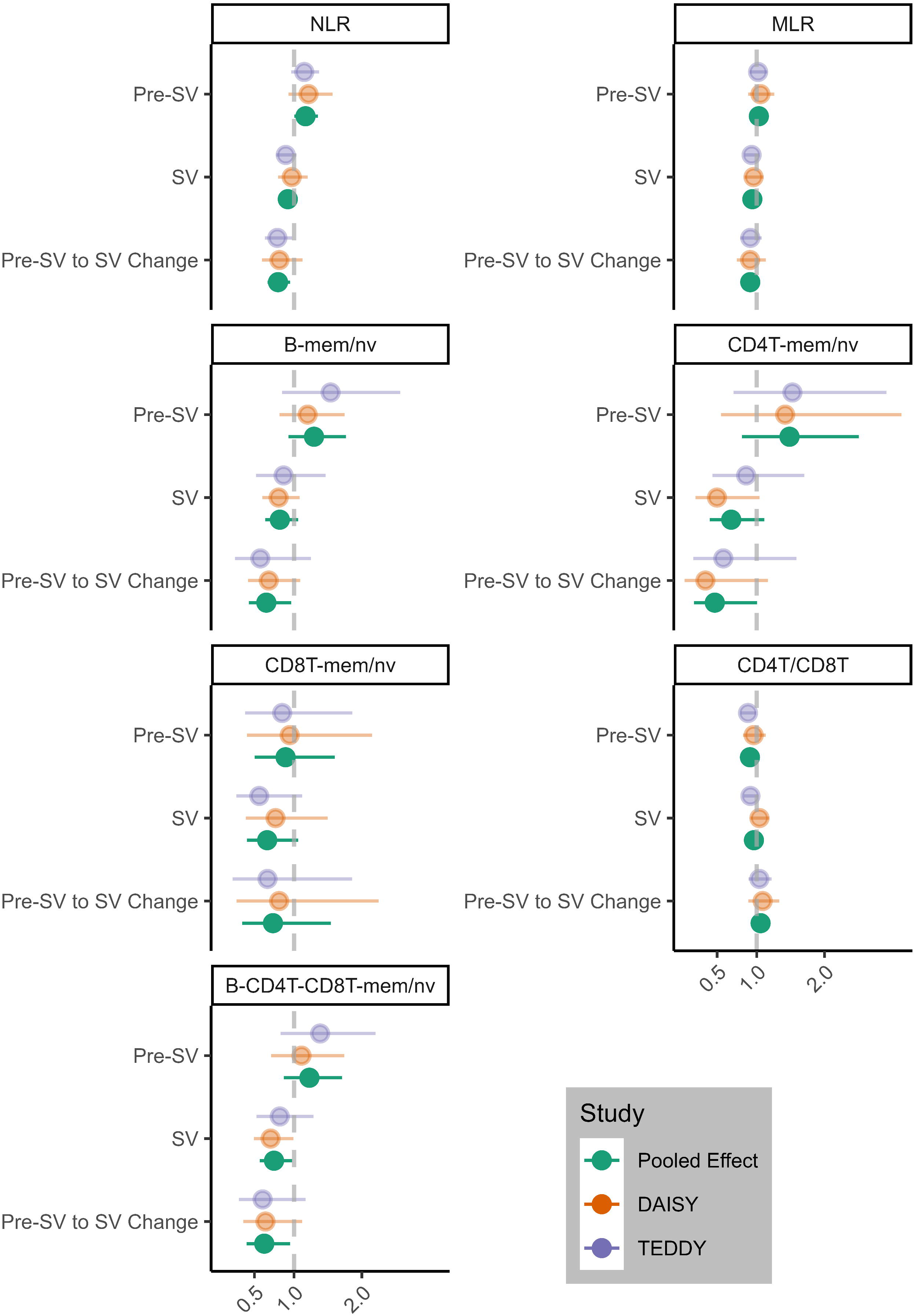
Meta-analysis: Differences in mean immune cell ratios between T1D cases and controls. * Abbreviations: NLR (neutrophil/lymphocyte ratio), MLR (monocyte/lymphocyte ratio), B-mem/nv (B-memory/B-naïve ratio), CD4T-mem/nv (CD4T-memory/ CD4T-naïve ratio), CD8T-mem/nv (CD8T-memory/CD8T-naïve ratio), CD4T/CD8T (CD4T/CD8T ratio), B-CD4T-CD8T-mem/nv (B-CD4T-CD8T-memory/B-CD4T-CD8T-naïve ratio), Pre-SV (pre-seroconversion), SV (seroconversion), T1D (type 1 diabetes). Interpretation: an immune cell ratio < 1 indicates a lower immune cell ratio among T1D cases compared to controls, and vice versa for an immune cell ratio > 1. In the pre-SV to SV change, an immune cell ratio < 1 indicates a reduced trajectory among T1D cases compared to controls, and vice versa for an immune cell ratio > 1.

**Figure 3.**
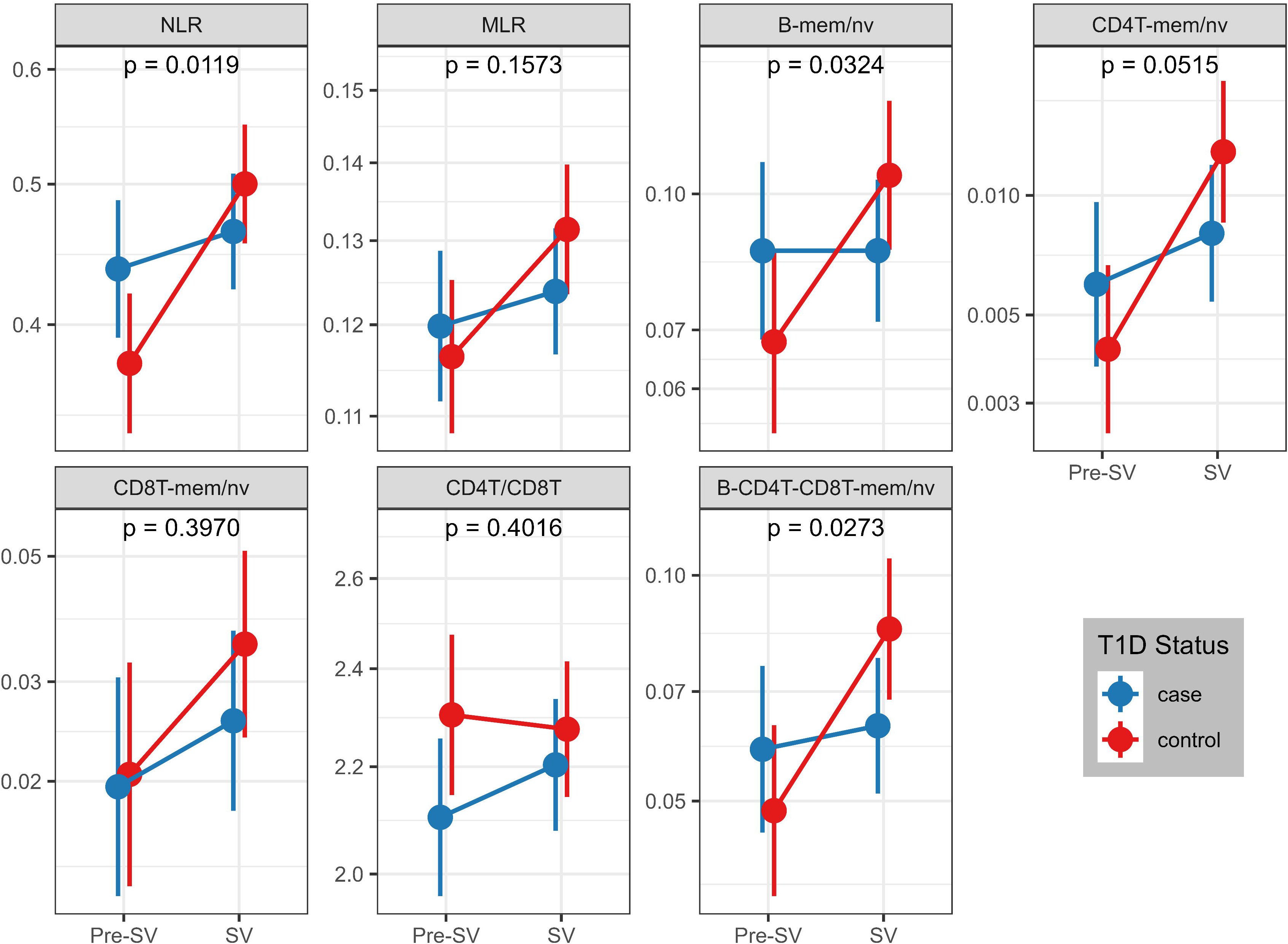
Immune cell ratios between T1D cases and controls around seroconversion. * Abbreviations: NLR (neutrophil/lymphocyte ratio), MLR (monocyte/lymphocyte ratio), B-mem/nv (B-memory/B-naïve ratio), CD4T-mem/nv (CD4T-memory/CD4T-naïve ratio), CD8T-mem/nv (CD8T-memory/CD8T-naïve ratio), CD4T/CD8T (CD4T/CD8T ratio), B-CD4T-CD8T-mem/nv (B-CD4T-CD8T-memory/B-CD4T-CD8T-naïve ratio), Pre-SV (pre-seroconversion), SV (seroconversion), T1D (type 1 diabetes). Interpretation: immune cell ratio changes between T1D cases and controls across the pre-SV and SV time points.

## Discussion

Among children who developed type 1 diabetes, we identified early and dynamic changes in immune cell ratios compared to controls. Some of these changes included evidence of increased innate immune activation, altered T cell balance, and delayed adaptive immune maturation during the period surrounding IA seroconversion. Specifically, we observed elevated NLR and reduced CD4T/CD8T cell ratios prior to seroconversion, followed by lower B-CD4T-CD8T-mem/nv ratios during seroconversion, and attenuated increases in the NLR, B-mem/nv, and B-CD4T-CD8T-mem/nv ratios over time. Since these immune cell ratios were estimated around the time of seroconversion to IA, they may provide insights into the etiology and pathophysiology of the type 1 diabetes. Given the natural changes in immune cell composition with age, having well-matched controls remained crucial for this study.

We found that type 1 diabetes cases started with higher NLR pre-SV, but they did not change as much pre-SV to SV as did controls. The NLR is considered a biomarker of systemic inflammation or activation of the innate immune response, and has seen previous use in cancer, infections, cardiovascular disease, and type 2 diabetes (T2D) in which a higher NLR typically underscored clinical worsening [7, 29–32]. After type 1 diabetes diagnosis, a higher NLR has been associated with diabetic ketoacidosis and carotid artery atherosclerosis [33, 34], while a lower NLR was associated with macroalbuminuria [35]. Salami et al. (2018) found reduced neutrophil counts among Swedish boys aged 4-12 years with islet autoantibodies in TEDDY [36]. However, our study included younger children prior to IA and analyzed neutrophil ratios rather than absolute counts, which might explain the differing effect direction. The elevated NLR at pre-SV in type 1 diabetes cases may be capturing an innate immune response to an environmental stimulus or otherwise detecting neutrophil activation, possibly contributing to autoimmunity. Thus, the NLR could have potential utility as an early marker of type 1 diabetes risk prior to IA. The reduced NLR change from pre-SV to SV in type 1 diabetes cases may suggest several physiological developments after the initial inflammatory response. Specifically, the decrease in circulating neutrophils could reflect pancreatic infiltration and contribution to insulitis; the increase in lymphocytes may indicate B and T cell recruitment as part of autoimmunity; alternatively, these shifts might signal a broader imbalance between innate and adaptive immune responses.

Type 1 diabetes cases had lower CD4T/CD8T ratios at pre-SV compared to controls. The ratio between CD4+ and CD8+ T cells has been identified as a marker of immune senescence and activation, as well as the balance between helper and cytotoxic T cell functions within the adaptive immune system. It has seen predominant use in studies pertaining to HIV and other infectious diseases [37, 38]. In the case of HIV, people with lower CD4T/CD8T ratios had reduced activity of CD4+ T helper cells and augmented activity of CD8+ cytotoxic T cells, reflective of their susceptibility to infection and ongoing viral replication [37]. In contrast, a higher CD4T/CD8T ratio was associated with acute respiratory distress syndrome and mortality during SARS-CoV-2 infection [38]; differences likely explained by HIV suppressing the immune response and SARS-CoV-2 activating it. For type 1 diabetes, previous work from Al-Sakkaf et al. (1989) observed reductions in the CD4T/CD8T ratio among participants who developed type 1 diabetes [9]. In adult onset type 1 diabetes, one study found decreased CD4+ and increased CD8+ T cell proportions, suggesting a similar relationship between helper and cytotoxic T cells [8]. Ferreira et al. (2010) uncovered a quantitative trait loci (QTL) located within the major histocompatibility complex for the CD4T/CD8T ratio, which was associated with class II variants in type 1 diabetes [39]. In our results, the decreased CD4T/CD8T ratio corroborates the previous work and reveals its potential role as an early indicator of disease risk prior to IA. This early shift in helper vs. cytotoxic T cells could suggest functional impairment of the CD4+ cells, including regulatory T cells, and subsequent activation of CD8+ cells into targeting pancreatic β-cells.

We found that type 1 diabetes cases had lower B-CD4T-CD8T-mem/nv ratio at SV and attenuated changes for B-mem/nv and B-CD4T-CD8T-mem/nv ratios compared to controls. B and T-cell memory/naïve ratios serve as relative measures of balance between the cells of the adaptive immune system, and similar inverted ratios (naïve/memory) have been investigated in diseases such as cancer, where higher relative ratios for CD4+ and CD8+ cells indicated poorer overall survival and a lower relative CD8+ naïve-to-memory ratio had longer progression-free survival [10]. In type 1 diabetes, the role of CD4+ and CD8+ T cells in autoimmunity is well established, though, B cells also take part via antibody secretion, antigen presentation, and pro-inflammatory cytokines [40]. Luo et al. (2025) found an increased level of naïve B lymphocytes among people with newly diagnosed type 1 diabetes [41]. People with adult onset type 1 diabetes also seemed to exhibit an increased level of naïve T cells and decreased level of effector memory subsets [8]. Our findings of a lower B-CD4T-CD8T mem/nv ratio at SV and reduced trajectories from pre-SV to SV for both B and B-CD4T-CD8T mem/nv ratios in type 1 diabetes cases compared to controls may signify an expansion or persistence of naïve cells. Or these patterns could reflect an impaired ability to activate into memory subsets following antigen exposure. In this way, people who go on to develop type 1 diabetes may possess an immature immunologic phenotype which predisposes them to autoimmunity and subsequent type 1 diabetes.

A major strength of our study includes longitudinal DNAm data prior to the onset of IA and thereafter, which allowed this novel look at immune cell profiles just before and during seroconversion. Meta-analysis of DAISY and TEDDY improved our power to detect these changes and establishes generalizability to high-risk children in Europe and the U.S. Use of the newest high-resolution cell deconvolution reference panel provides greater granularity over the common six-cell panel, including allowing us to distinguish between memory and naïve cell populations, albeit through indirect measures of immune cell composition.

Our study also possessed some limitations. Since we did not have access to direct flow cytometry data, we utilized cell deconvolution of peripheral blood samples for immune cell estimates with a reference panel derived from healthy adult donors, which included both sexes and several ethnic and ancestral backgrounds. Despite the unavailability of flow cytometry data, we leveraged DNAm data to estimate these proportions with a highly accurate, reference-based immune profiling method that has been previously used for similar tasks in cancer, cardiovascular disease, and neurological disease [42–44]. Regardless of the difference in age for the reference, our immune cell proportions seemed to correspond to previously reported reference ranges for this age group [24]. Despite differences in age at type 1 diabetes and IA onset across DAISY and TEDDY, immune profiles changes were robust across studies, independent of age. Future research should consider the creation and utilization of a peripheral blood reference panel for DNAm data specific to children and adolescents, since immunological development and subsequent changes occur with age.

In this epidemiologic analysis leveraging two well-characterized, prospective cohorts, we identified distinct alterations in immune cell ratios that precede and accompany IA in children who progress to type 1 diabetes. The findings herein suggest potential roles for the NLR and CD4T/CD8T ratio in serving as early biomarkers for type 1 diabetes risk stratification, even before the onset of IA. Having the ability to identify these high-risk individuals provides a unique opportunity for public health intervention. In addition, children who develop type 1 diabetes appeared to have lower B-CD4T-CD8T mem/nv ratios and decreased trajectories for the NLR, B mem/nv, and B-CD4T-CD8T mem/nv ratios and may serve as indicators of immune dysfunction or immaturity and type 1 diabetes progression. Together, these findings underscore the importance of temporal dynamics in immune cell profiles as novel biomarkers of early disease progression and offer potential targets for improved risk stratification and timely intervention aimed at delaying or preventing type 1 diabetes. Moreover, high-resolution cell deconvolution represents a cost-effective, scalable approach for observational studies with DNAm data to profile immunodynamics specific to disease in the absence of flow cytometry. Future work will need to verify these results through functional analyses to determine the biological relevance of the variations in the DNAm-derived immune cell proportions and immune cell ratios.

## Acknowledgments

We would like to acknowledge the dedication of the study participants in DAISY and TEDDY and their families, without which these studies would not be possible, as well as the assiduous effort of the research and data coordination staff.

## Author Contributions (CRediT – Contributor Role Taxonomy)

KRH: Conceptualization, Formal analysis, Methodology, Software, Validation, Visualization, Writing - Original Draft, Writing - Review & Editing

PMC: Conceptualization, Methodology, Writing - Review & Editing

SDS: Data Curation, Writing - Review & Editing

CTR: Data Curation, Writing - Review & Editing

LAV: Software, Writing - Review & Editing

KK: Writing - Review & Editing

MR: Funding acquisition, Investigation, Project administration, Resources, Writing - Review & Editing

JMN: Funding acquisition, Investigation, Project administration, Writing - Review & Editing

RKJ: Conceptualization, Funding acquisition, Methodology, Validation, Resources, Supervision, Writing - Review & Editing

## Guarantor Statement

R.K.J. is the guarantor of this work and, as such, had full access to all the data in the study and takes responsibility for the integrity of the data and the accuracy of the data analysis.

## Conflict of Interest

The authors declare that they have no conflicts of interest related to the study.

## Funding

DAISY was funded by R01DK032493 (to MR) and R01DK104351 (to JMN). This work was supported by Leona M. and Harry B. Helmsley Charitable Trust (2103-05094) to JMN. This research was performed using resources generated by the TEDDY study group, a collaborative clinical study sponsored by the National Institute of Diabetes and Digestive and Kidney Diseases (NIDDK), National Institute of Allergy and Infectious Diseases (NIAID), National Institute of Child Health and Human Development (NICHD), National Institute of Environmental Health Sciences (NIEHS), Centers for Disease Control and Prevention (CDC), and Breakthrough T1D (formerly JDRF). This manuscript was not prepared under the auspices of the TEDDY study group and does not necessarily reflect the opinions and views of the TEDDY study, the CDC, or NIH.

## Prior Presentation information

A previous version of this work was presented as a poster at the Annual Meeting of the Society for Epidemiologic Research in 2024.

## Notes

### Competing Interest Statement

The authors have declared no competing interest.

### Summary of Updates

Funding for the study was updated to match language from the data coordinating center for TEDDY.

