## Supplemental Material for "Differences in immune cell profiles around the time of islet autoimmunity seroconversion in children with and without type 1 diabetes"

Supplemental Table 1. Comparison of immune cell subsets by age in TEDDY to reference ranges

|  | Samples | TEDDY (age 1 - 2 years) | Reference (age 1-2 years)^a^ |
| --- | --- | --- | --- |
| Immune subset | N | Median (10th, 90th perc.) |  |
| B cell | 395 | 15 (11-21) | 25 (16-35) |
| Natural killer cell | 395 | 5 (02-9) | 7 (03-15) |
| Helper T cell | 395 | 25 (17-33) | 41 (32-51) |
| Cytotoxic T cell | 395 | 12 (7-17) | 20 (14-30) |

a. Reference data extracted from Shearer et al. 2003, where Immune subset percentages were based on flow cytometry that gated on cells with a CD45 anchor marker.

Supplemental Figure 2. Immune cell proportions in TEDDY participants across time

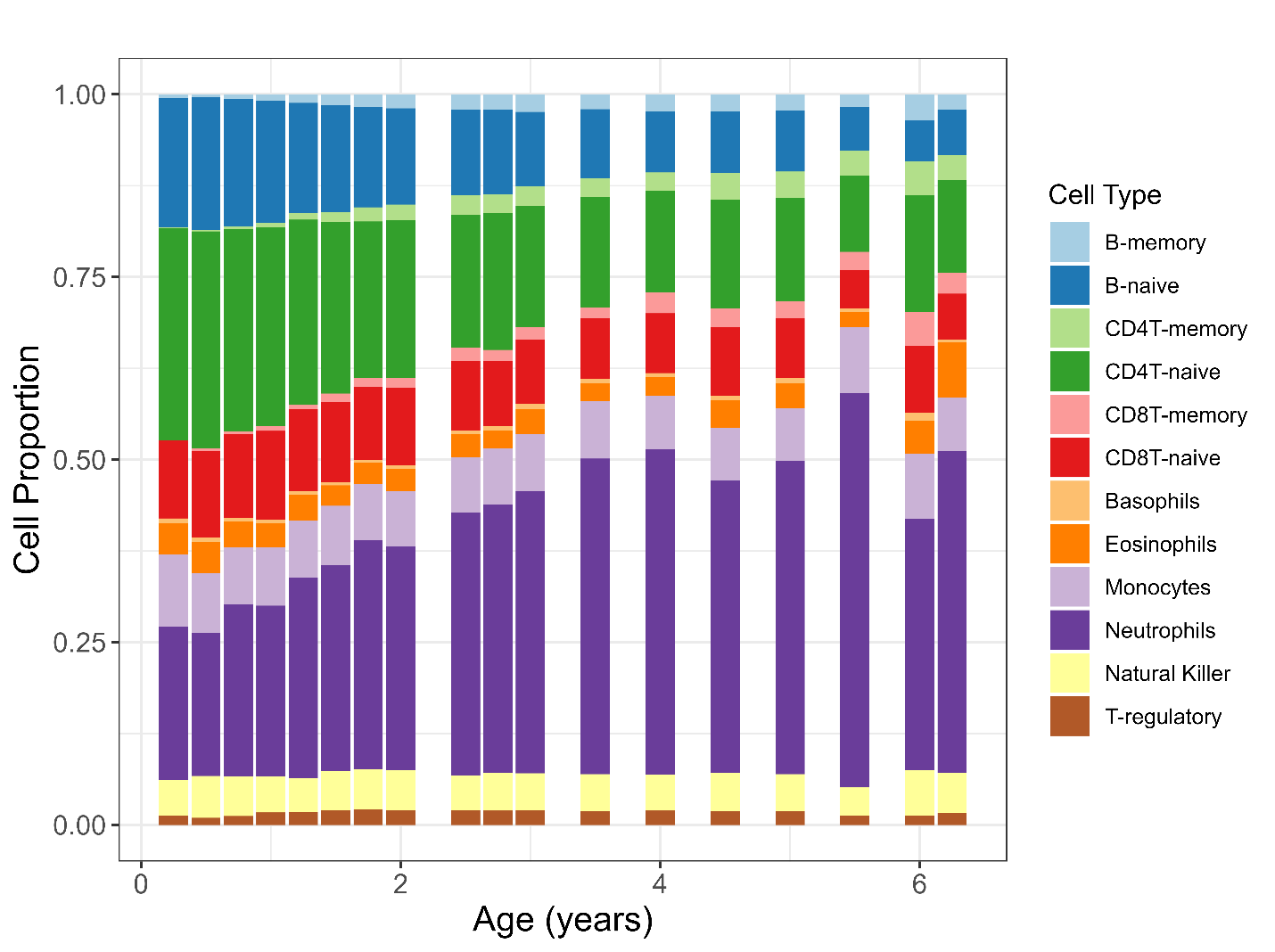

* Includes all TEDDY samples that passed quality control checks and study exclusions.

Supplemental Appendix 1. Additional information on sample pre-processing in DAISY

For each pair of technical replicates, we retained the sample that matched the platform used for the majority of that participant's samples. If an equal number of samples were available on each platform, the sample measured on the EPIC array was prioritized. To minimize potential platform-related batch effects, we prioritized selecting samples that captured the SV timepoint of interest, ensuring that all retained samples for a given participant came from the same array platform.
